# Preclinical evaluation of Imatinib does not support its use as an antiviral drug against SARS-CoV-2

**DOI:** 10.1101/2020.11.17.386904

**Authors:** Franck Touret, Jean-Sélim Driouich, Maxime Cochin, Paul Rémi Petit, Magali Gilles, Karine Barthélémy, Grégory Moureau, Francois-Xavier Mahon, Denis Malvy, Caroline Solas, Xavier de Lamballerie, Antoine Nougairède

## Abstract

Following the emergence of SARS-CoV-2, the search for an effective and rapidly available treatment was initiated worldwide based on repurposing of available drugs. Previous reports described the antiviral activity of certain tyrosine kinase inhibitors (TKIs) targeting the Abelson kinase 2 against pathogenic coronaviruses. Imatinib, one of them, has more than twenty years of safe utilization for the treatment of hematological malignancies. In this context, Imatinib was rapidly evaluated in clinical trials against Covid-19. Here, we present the pre-clinical evaluation of Imatinib in multiple models. Our results indicated that Imatinib and another TKI, the Masitinib, exhibit an antiviral activity in VeroE6 cells. However, Imatinib was inactive in a reconstructed bronchial human airway epithelium model. *In vivo*, Imatinib therapy failed to impair SARS-CoV-2 replication in a golden Syrian hamster model despite high concentrations in plasma and in the lung. Overall, these results do not support the use of Imatinib and similar TKIs as antivirals in the treatment of Covid-19.

## Introduction

Coronaviruses are enveloped positive-stranded RNA viruses belonging to the order *Nidovirales*, which have a long history of causing respiratory tract infections. During the 21^st^ century, the emergence of the severe acute respiratory syndrome coronavirus (SARS-CoV) and the Middle East respiratory syndrome coronavirus (MERS-CoV), both associated with a high proportion of severe acute respiratory syndromes, raised fears of coronavirus-induced pandemic (de Wit et al., 2016). In late 2019, another coronavirus, named severe acute respiratory syndrome coronavirus-2 (SARS-CoV-2) emerged in China. Once again this new viral pathogen was associated with severe respiratory infections (Zhu et al., 2020). SARS-CoV-2 human disease, called coronavirus disease 2019 (Covid-19), was characterized as a pandemic on March 2020 by the World Health Organization. To date, there is no licensed vaccine against SARS-CoV-2 but many candidates are in development (Jeyanathan et al., 2020). Without licensed antiviral drugs that specifically target coronaviruses, drug repurposing has been considered as the fastest strategy to find an antiviral specific therapy since the conventional development of new drugs may take several years (Manganaro et al., 2020). However, and after a plethora of clinical trials in a desperate attempt to rapidly find efficient antiviral therapies and adjunctive treatments for the management of moderate to critical Covid-19, the only breakthrough, yet, was a supportive therapy (The RECOVERY Collaborative Group, 2020).

A few years ago, a screen of clinically developed drugs identified two tyrosine kinase inhibitors (TKIs), Imatinib and Dasatinib as potential inhibitors of SARS-CoV and MERS-CoV replication *in vitro* (Dyall et al., 2014). In VeroE6 cells, Imatinib exhibited 50% dose effective concentrations (EC50) of 17.7µM and 9.8µM against MERS-CoV and SARS-CoV, respectively. Imatinib antiviral activity was then more precisely investigated *in vitro*, in multiple cell lines and with three different coronavirus: SARS-CoV, MERS-CoV and infectious bronchitis virus (IBV) (Coleman et al., 2016; Sisk et al., 2018). Hence, imatinib inhibits coronavirus replication cycle prior to viral RNA replication probably by preventing the coronavirus spike protein mediated cell fusion (Sisk et al., 2018). This blockage is the consequence of the inhibition of a specific tyrosine kinase, the Abelson tyrosine-protein kinase 2 (Abl2)(Coleman et al., 2016). Indeed, some TKIs, such as the Imatinib, target multiple tyrosine kinases including the Abelson tyrosine-protein kinase 1 and 2 (Abl1 and 2), the stem cell factor receptor (c-KIT/CD117) and the platelet-derived growth factor receptor (PDGF-R). In the context of the current Covid-19 pandemic, these reports took a new twist and opened the door to the direct availability of potential direct antiviral drugs against SARS-CoV-2. Indeed Imatinib, which has been initially developed twenty years ago for the treatment of chronic myeloid leukaemia (CML), has a long history of human use (Henkes et al., 2008). Consequently, several clinical trials using Imatinib were started in Covid-19 patients (Ashkan, 2020; ROUSSELOT, 2020). However, the antiviral activity of TKIs against SARS-CoV-2 was not thoroughly evaluated *in vitro* and *in vivo*.

Here we present the preclinical evaluation of tyrosine kinase inhibitors, especially the Imatinib, against SARS-CoV-2. First, we have evaluated the efficacy of three kinase inhibitors *in vitro* and we then assessed the Imatinib antiviral activity using bronchial human airway epithelium (HAE) and *in vivo*.

## Material and methods

### Cell line

VeroE6 (ATCC CRL-1586) cells were grown in minimal essential medium (Life Technologies) with 7⍰5% heat-inactivated fetal calf serum (FCS; Life Technologies), at 37°C with 5% CO_2_ with 1% penicillin/streptomycin (PS, 5000U.mL^−1^ and 5000µg.mL^−1^ respectively; Life Technologies) and supplemented with 1⍰% non-essential amino acids (Life Technologies).

### Human airway epithelia (HAE)

Mucilair™ HAE reconstituted from human primary cells of bronchial biopsies were purchased from Epithelix SARL (Geneva, Switzerland). The bronchial epithelium was maintained in air-liquid interface with specific media purchased from Epithelix. For the first HAE experiment, epithelium derived from a 56-year-old donor Caucasian female with no existing pathologies reported or detected. The second and third set of HAE experiment, epithelium derived from a 17-year-old donor Hispanic male with no existing pathologies reported or detected. All samples purchased from Epithelix SARL have been obtained with informed consent. These studies were conducted according to the declaration of Helsinki on biomedical research (Hong Kong amendment, 1989), and received approval from local ethics committee.

### Virus strain

SARS-CoV-2 strain BavPat1 was obtained from Pr. C. Drosten through EVA GLOBAL (https://www.european-virus-archive.com/). To prepare the virus working stock, a 25cm^2^ culture flask of confluent VeroE6 cells growing with MEM medium with 2.5% FCS was inoculated at MOI 0.001. Cell supernatant medium was harvested at the peak of replication and supplemented with 25mM HEPES (Sigma-Aldrich) before being stored frozen in aliquots at -80°C. All experiments with infectious virus were conducted in a biosafety level 3 laboratory.

### EC50 and CC50 determination

One day prior to infection, 5×10^4^ VeroE6 cells per well were seeded in 100µL assay medium (containing 2.5% FCS) in 96 well culture plates. The next day, eight 2-fold serial dilutions of compounds (from 0.62 µM to 40µM for Imatinib/Asciminib and from 0.78µM to 100µM for Masitinib) in triplicate were added to the cells (25µL/well, in assay medium). Four virus control wells were supplemented with 25µL of assay medium. After 15 min, 25µL of a virus mix diluted in medium was added to the wells. The amount of virus working stock used was calibrated prior to the assay, based on a replication kinetics, so that the viral replication was still in the exponential growth phase for the readout as previously described (Delang et al., 2016; Touret et al., 2020, 2019). Four cell control wells (*i*.*e*. with no virus) were supplemented with 50µL of assay medium. On each culture plate, a control compound (Remdesivir, BLDpharm) was added in duplicate with eight 2-fold serial dilutions (0.16µM to 20µM). Plates were incubated for 2 days at 37°C prior to quantification of the viral genome by real-time RT-PCR. To do so, 100µL of viral supernatant was collected in S-Block (Qiagen) previously loaded with VXL lysis buffer containing proteinase K and RNA carrier. RNA extraction was performed using the Qiacube HT automat and the QIAamp 96 DNA kit HT following manufacturer instructions. Viral RNA was quantified by real-time RT-qPCR (GoTaq 1-step qRt-PCR, Promega) using 3.8µL of extracted RNA and 6.2µL of RT-qPCR mix and standard fast cycling parameters, *i*.*e*., 10min at 50°C, 2 min at 95°C, and 40 amplification cycles (95°C for 3 sec followed by 30sec at 60°C). Quantification was provided by four 2 log serial dilutions of an appropriate T7-generated synthetic RNA standard of known quantities (10^2^ to 10^8^ copies/reaction). RT-qPCR reactions were performed on QuantStudio 12K Flex Real-Time PCR System (Applied Biosystems) and analyzed using QuantStudio 12K Flex Applied Biosystems software v1.2.3. Primers and probe sequences, which target SARS-CoV-2 N gene, were: Fw: GGCCGCAAATTGCACAAT; Rev: CCAATGCGCGACATTCC; Probe: FAM-CCCCCAGCGCTTCAGCGTTCT-BHQ1. The 50% and 90% effective concentrations (EC50, EC90; compound concentration required to inhibit viral RNA replication by 50% and 90%) were determined using logarithmic interpolation as previously described (Touret et al., 2019). For the evaluation of the 50% cytotoxic concentrations (CC50), the same culture conditions as for the determination of the EC50 were used, without addition of the virus, and cell viability was measured using CellTiter Blue® (Promega) following manufacturer’s instructions. CC50 was determined using logarithmic interpolation. All data obtained were analyzed using GraphPad Prism 7 software (Graphpad software).

### Antiviral assay using HAE

After a gentle wash with pre-warmed OPTI-MEM medium (Life technologies), epithelia were infected with SARS-COV-2 on the apical side using a MOI of 0.1 as previously described (Pizzorno et al., 2020). Cells were cultivated in basolateral media that contained different Imatinib concentrations or without the drug (Virus control). Remdesivir (BLD pharm) was used as a positive drug control. Media were renewed every day during the experiment. At day 1, before the media renewing the apical side of the epithelia was washed with warm OPTI-MEM in order to eliminate the viral inoculum. Samples were collected at the apical side by washing with 200µL of pre-warmed OptiMEM medium: 100µL was used for RNA extraction using the Qiacube HT and viral RNA were quantified as described in the ‘EC50 and CC50 determination’ section; the remaining quantity was used to perform a TCID_50_ assay. At day 4 post-infection, total intracellular RNA of each well was extracted using RNeasy 96 HT kit (Qiagen) following manufacturer’s instructions and viral RNA were quantified as described in the ‘EC50 and CC50 determination’ section.

### *In vivo* experiments

*In vivo* experiments were approved by the local ethical committee (C2EA—14) and the French ‘Ministère de l’Enseignement Supérieur, de la Recherche et de l’Innovation’ (APAFIS#23975) and performed in accordance with the French national guidelines and the European legislation covering the use of animals for scientific purposes. All experiments were conducted in BSL 3 laboratory.

#### Animal handling

Three-week-old female golden Syrian hamsters were provided by Janvier Labs. Animals were maintained in ISOcage P - Bioexclusion System (Techniplast) with unlimited access to water/food and 14h/10h light/dark cycle. Animals were weighed and monitored daily for the duration of the study to detect the appearance of any clinical signs of illness/suffering.

#### Infection

Virus inoculation was performed under general anesthesia (isoflurane). Four-week-old anesthetized animals were intranasally infected with 50µL containing 10^4^ TCID_50_ of virus in 0.9% sodium chloride solution). The mock-infected group was intranasally inoculated with 50µL of 0.9% sodium chloride solution.

#### Drug administration

One 400mg tablet of Imatinib mesylate was crushed and Imatinib solubilized in 25mL of distilled water buffered to pH=4.6 using an aceto-acetic buffer in order to obtain a concentration of 16 mg/mL (32.41mM). Concentration was confirmed by liquid chromatography coupled with mass spectrometry before animal administration (see below). Twice a day, hamsters were orally treated with different doses of Imatinib or with a 0.9% sodium chloride solution (control group), under general anesthesia (isoflurane). For control groups treated with Favipiravir (see below), hamsters were intraperitoneally treated.

#### Lungs and blood collection

Lung and blood samples were collected immediately after the time of sacrifice at the end of the dosing interval. The left pulmonary lobe was first washed in 10mL of 0.9% sodium chloride solution, blotted with filter paper, weighed and then transferred to a 2mL tube containing 1mL of 0.9% sodium chloride solution and 3mm glass beads. They were crushed using a Tissue Lyser machine (Retsch MM400) for 20min at 30 cycles/s and then centrifuged 10min à 16,200g. Supernatant media were transferred to 1.5mL tube, centrifuged 10 min at 16,200g and stored at -80°C. One milliliter of blood was harvested in a 2mL tube containing 100µL of 0.5M EDTA (ThermoFischer Scientific). Quantitative real-time RT-PCR and TCID_50_ assays were performed as described below.

#### Quantitative real-time RT-PCR (RT-qPCR) assays

To avoid contamination, all experiments were conducted in a molecular biology laboratory that was specifically designed for clinical diagnosis, and which includes separate laboratories for each step of the procedure. Prior to PCR amplification, RNA extraction was performed using the QIAamp 96 DNA kit and the Qiacube HT kit and the Qiacube HT (both from Qiagen) following the manufacturer’s instructions. Shortly, 100 μL of plasma or organ clarified homogenates was spiked with 10μL of internal control (bacteriophage MS2), and then transferred into a S-block containing the recommended volumes of VXL, proteinase K and RNA carrier. RT-qPCR (SARS-CoV-2 and MS2 viral genome detection) were performed with the Express one step RT-qPCR Universal kit (EXPRESS One-Step Superscript™ qRT-PCR Kit, universal Invitrogen) using 3.5µL of RNA and 6.5µL of RT qPCR mix and standard fast cycling parameters, *i*.*e*., 10min at 50°C, 2 min at 95°C, and 40 amplification cycles (95°C for 3 sec followed by 30sec at 60°C). Quantification was provided by four 2 log serial dilutions of an appropriate T7-generated synthetic RNA standard of known quantities (10^2^ to 10^8^ copies/reaction). RT-qPCR reactions were performed on QuantStudio 12K Flex Real-Time PCR System (Applied Biosystems) and analyzed using QuantStudio 12K Flex Applied Biosystems software v1.2.3. Primers and probes sequences used to detect SARS-CoV-2: RNAdependent RNA polymerase Fwd: 5’-GTGARATGGTCATGTGTGGCGG-3’ Rev: 5’-CARATGTTAAASACACTATTAGCATA-3’ Probe: 5’-FAM-CAGGTGGAACCTCATCAGGAGATGC-TAMRA-3 (Corman et al., 2020). Bacteriophage MS2 primers: Fwd: 5’-CTCTGAGAGCGGCTCTATTGGT-3’ Rev: 5’-GTTCCCTACAACGAGCCTAAATTC-3’ Probe: 5’-VIC-TCAGACACGCGGTCCGCTATAACGA-TAMRA-3’ (Ninove et al., 2011).

### Tissue-culture infectious dose 50 (TCID_50_) assay

To determine infectious titers, 96-well culture plates containing confluent VeroE6 cells were inoculated with 150μL per well of serial dilutions of each sample (four-fold or ten-fold dilutions when analyzing lung clarified homogenates or cell supernatant media respectively). Each dilution was analyzed in sextuplicate. Plates were incubated for 4 days and then read for the absence or presence of cytopathic effect in each well. Infectious titers were estimated using the method described by Reed & Muench (Reed and Muench, 1938) and expressed as TCID_50_/ml or TCID_50_/g of lung.

### Imatinib quantification in plasma and lung tissues

Quantification of Imatinib in plasma and lung tissues (collected as described above) was performed by a validated sensitive and selective validated high-performance liquid chromatography coupled with tandem mass spectrometry method (UPLC-TQD, Waters, USA) as previously described (Ferrer et al., 2020). Imatinib was extracted by a simple protein precipitation, using acetonitrile for plasma and ice-cold acetonitrile for clarified lung homogenates, containing the isotopic internal standard (Imatinib-d8, Alsachim). The method was validated according to the EMA guidelines (European Medicines Agency, 2018) with a lower limit of quantification of 25 ng/mL (precision: 5.42%; accuracy: 3.87%). Precision and accuracy of the three quality control samples (QCs) were within 15% over the calibration range (25 ng/mL to 5000 ng/mL).

To assess the selectivity and specificity of the method and matrix effect applied to animal study, blank plasma and tissues homogenates from two control animals (uninfected and untreated) were processed at each run. Moreover, the same control samples spiked with Imatinib concentration equivalent to QCs (100, 500 and 3000 ng/mL) were also processed and compared to the QCs samples. Ten-fold dilution integrity was validated with both precision and accuracy within 15% to allow quantification of the highest doses evaluated.

### Statistical analysis

Graphical representations and statistical analyses were performed with Graphpad Prism 7 (Graphpad software). Statistical details for each experiments are described in the figure legends and in corresponding supplemental tables. When relevant, two-sided statistical tests were always used. P-values lower than 0.05 were considered statistically significant.

## Results

### Imatinib and Masitinib inhibit SARS-Cov-2 replication in VeroE6 cells

First, we evaluated the antiviral activity of several TKIs (Imatinib, Masitinib and Asciminib) against SARS-CoV-2 using an antiviral assay in VeroE6 cells as previously described (Touret et al., 2020). This assay estimates the viral inhibition by measuring the viral genomic RNA replication by qRT-PCR (Touret et al., 2019). Imatinib and Masinitib inhibited SARS-Cov-2 replication with EC50s of 2.5 and 2.3µM respectively (Fig 1 A, B and D), similar with recent reports for SARS-CoV-2 (Drayman et al., 2020; Han et al., 2020) but slightly lower than those found for SARS-CoV and MERS-CoV (Dyall et al., 2014). Imatinib was less cytotoxic than Masitinib resulting in a superior selectivity Index (SI=CC50/EC50): 16.2 versus 11.5 (Fig 1 A, B and D). Asciminib did not show any antiviral activity as its EC50 is similar to its CC50 resulting in a SI of 1.1 (Fig1 C, D).

**Figure 1:**
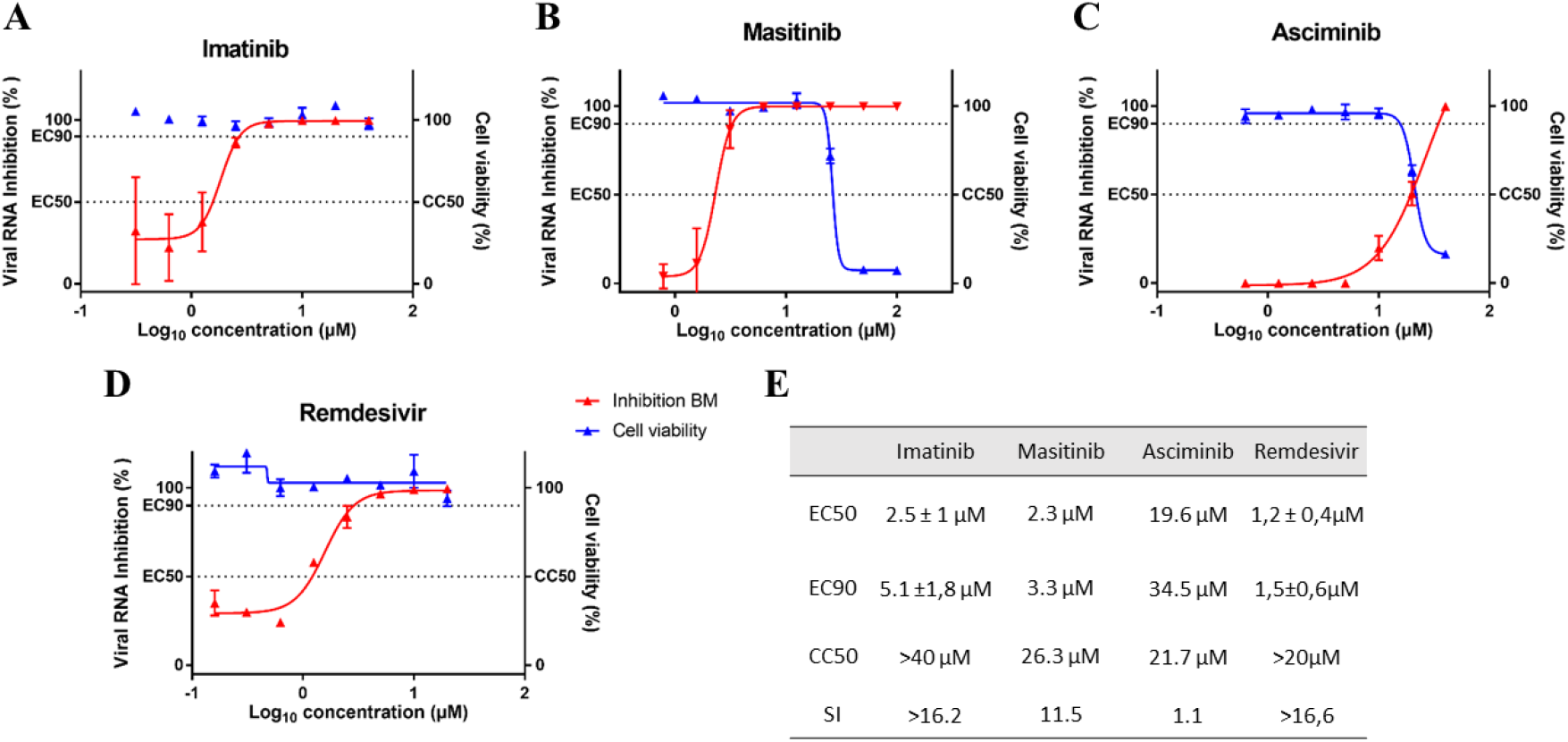
Antiviral activity of different tyrosine Kinase inhibitors in VeroE6 cells. Dose response curve and cell viability for Imatinib (A), Masitinib (B) and Asciminib (C) and control compound Remdesivir (D). E: Table of EC50, EC90, CC50 and SI of the different tyrosine kinase inhibitors. Results presented in the table for Imatinib and remdesivir are the mean±SD from three independent experiments.

### Imatinib is not active in a bronchial human airway epithelium (HAE)

To further characterize Imatinib antiviral activity we used a recently described model of reconstituted human airway epithelial of bronchial origin (Pizzorno et al., 2020) by testing in duplicate five different concentrations of Imatinib (10; 5; 2.5; 1.25; 0.6 µM). Epithelia were exposed to the drug during three days through their basolateral side (Fig 2A). Remdesivir at 10 µM was used as a positive control for antiviral activity (Pizzorno et al., 2020). We monitored viral excretion at the apical side of the epithelium from 2 dpi to 4 dpi by measuring viral RNA yields (Fig.3B) and infectious titres (Fig 2C) in apical washes using respectively quantitative real time RT-PCR and TCID_50_ assays. Despite a small non-significant reduction of viral RNA yields at 2 dpi with the highest concentration of Imatinib, results showed no significant antiviral activity at any dose. Similar results were found when measuring intracellular viral RNA yields at 4 dpi (Supplemental Fig1). In contrast, Remdesivir induced strong antiviral activity. In order to confirm these first results, the experiment was repeated two times and the same results were obtained: Imatinib treatment at the three highest doses had no impact on viral replication (Supplemental Fig2).

**Figure 2:**
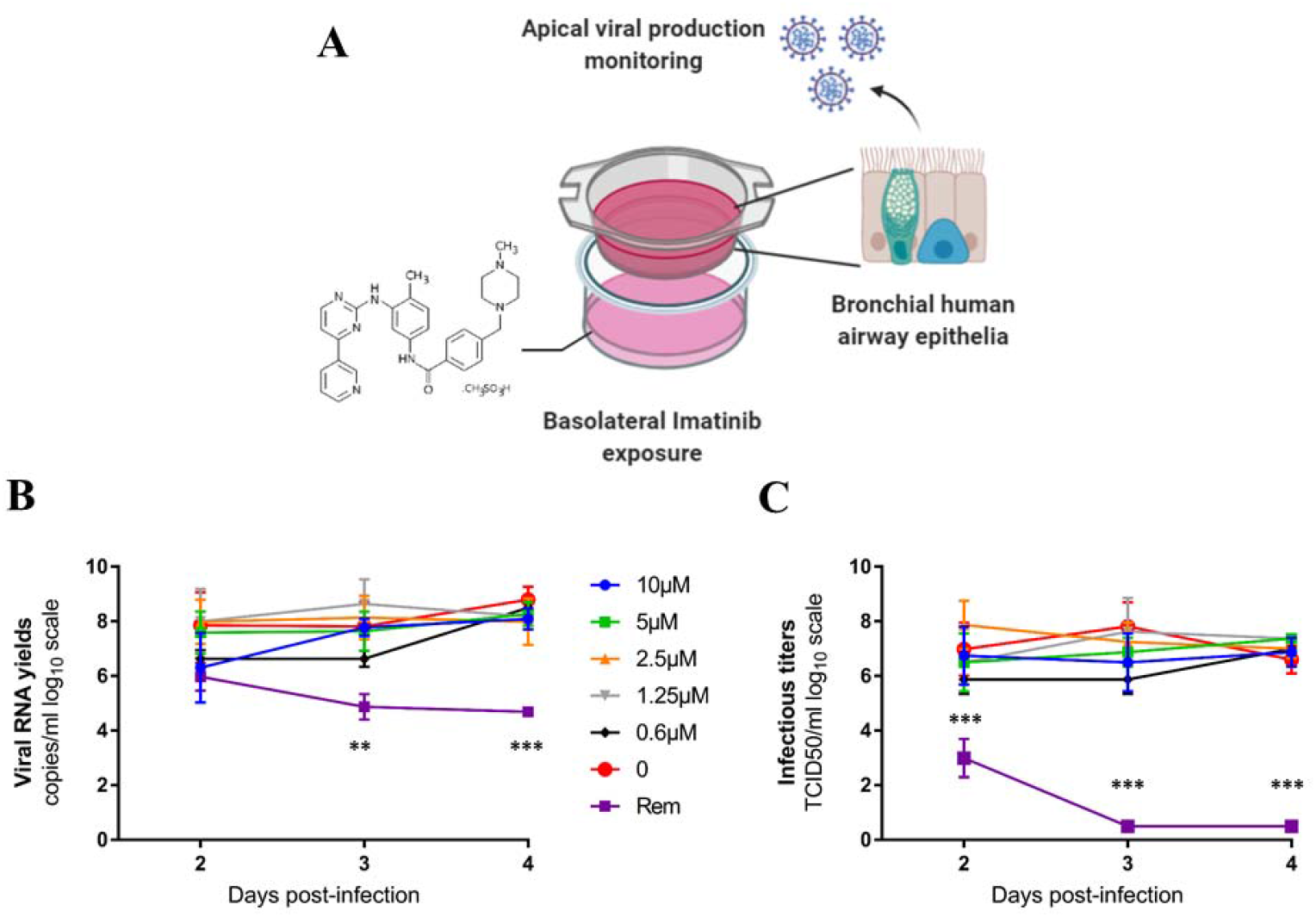
Antiviral activity of Imatinib in a bronchial human airway epithelium. Graphical representation of the experiment (A). Kinetics of virus excretion at the apical side of the epithelium measured using an RT-qPCR assay (B) and a TCID_50_ assay (C). Data represent mean±SD. Statistical significance was calculated by 1-way ANOVA versus untreated group. No statistical difference were observed with Imatinib regardless the drug concentration. Rem (Remdesivir at 10µM) was used as a positive drug control. ** and *** symbols indicate that the average value for the Rem group is significantly lower than that of the untreated group with a p-value ranging between 0.001-0.01 and 0.0001-0.001 respectively. The graphical representation was created with BioRENDER.

### Imatinib does not block SARS-CoV-2 replication in lungs of infected hamsters

As Imatinib had the best SI and was under evaluation in clinical trials, we evaluated its antiviral activity in a previously described hamster model of SARS-CoV-2 infection (Chan et al., 2020; Driouich et al., 2020). To assess the efficacy of Imatinib, groups of 4 hamsters were intranasally infected with 10^4^ TCID_50_ of SARS-CoV-2 and received the drug orally twice a day (BID). We used two doses of Imatinib: 16 and 32mg/day BID (corresponding to 267±23 and 514±45mg/kg/day BID respectively). A group of 4 hamsters that orally received saline solution BID was used as control. The treatment was initiated the day of infection and ended 2 days post-infection (dpi) (Fig 3A). Viral replication in lungs and plasma was assessed at 3 dpi. This experimental design, with animals infected with a low dose of SARS-CoV-2 and sacrificed at the peak of replication previously proved to be highly sensitive to assess the efficacy of Favipiravir (Driouich et al., 2020). In all experiments, Favipiravir was used as a positive control (50mg/day BID) as previously described (Driouich et al., 2020).

**Figure 3:**
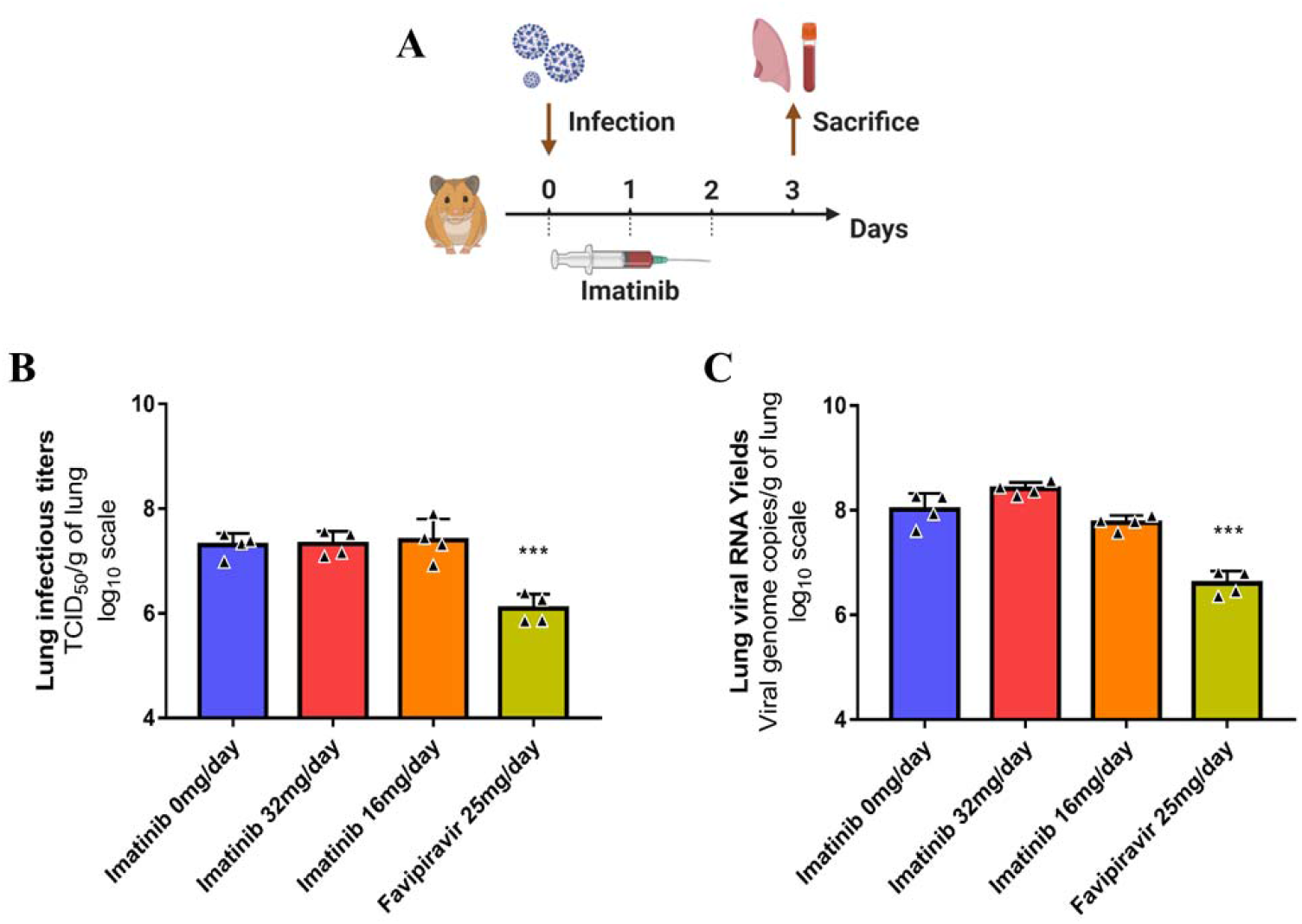
Antiviral activity of Imatinib in a Golden Syrian hamster model. A: Experimental timeline. Groups of 4 hamsters were intranasally infected with 10^4^ TCID_50_ of SARS-CoV-2 and received the Imatinib, orally twice a day. Favipiravir was used as a positive control. B: Lung infectious titers (measured using a TCID_50_ assay) expressed in TCID_50_/g of lung. C: Viral RNA yields (measured using an RT-qPCR assay) expressed in viral genome copies/g of lung. Data represent mean±SD. *** symbols indicate that the average value for the group is significantly lower than that of the untreated group with a p-value ranging between 0.0001-0.001. The experimental timeline was created with BioRENDER.

When analysis of virus replication in clarified lung homogenates was based on infectious titers (measured using TCID_50_ assay) or on viral RNA yields (measured using quantitative real time RT-PCR assay), no significant differences between treated and untreated animals were observed whatever the dose of Imatinib used (*p*≥0.7038 and *p*≥0.0558 respectively) (Fig 3B and C). In contrast, administration of Favipiravir led to significant reductions of both infectious titers and viral RNA yields in clarified lung homogenates (*p*=0.0005 and *p*=0.0003 respectively). Likewise, treatment with Imatinib, whatever the dose administrated, did not affect significantly the molecular plasma viral loads. Animals treated with the dose of 32mg/day BID of Imatinib showed signs of toxicity from 3 dpi, with their mean normalized weight becoming significantly lower than that of untreated animals (Supplemental Fig.3).

In order to confirm these first results, the experiment was repeated with groups of 6 hamsters and the same conclusions were made: administration of Imatinib at both dosing regimens had no impact on viral replication in clarified lung homogenates and plasma whereas Favipiravir therapy led to a significant reduction of viral replication in all conditions (Supplemental Fig4).

To assess the exposure to the drug during the first set of experiments, plasma and lung concentrations of Imatinib from sacrificed animals (*i*.*e*. trough concentrations after multiple administration) were determined (Table 1). Results showed a moderate inter-individual variability in plasma and lung concentrations (coefficient of variation from 26 to 58 %). A good penetration of Imatinib in lungs was observed with a plasma/lung percentage ratio ranging from 53 to 99% according to the dosing regimen. At both dosing regimens, lung concentrations of Imatinib were above the EC50. The inhibitory quotient (IQ: ratio between the concentration and the EC50) in plasma and lungs was 23 and 11 at 32 mg/day and 3 and 2.8 at 16 mg/day, respectively (table 1).

**Table 1:** Plasma and lung concentrations of Imatinib after administration of multiple doses

| Dosing regimen | Plasma (PI)* | Lung (L)* | ratio L/PI |
| --- | --- | --- | --- |
| **Imatinib**<br>**32mg/day BID** | 31.7 ± 8.2 µg/ml<br>(64,3 ± 16,8 µM) | 15.5 ± 5.7 µg/g<br>(31,4 ± 11,6 µM) | 52 ± 25 % |
| **Imatinib**<br>**16mg/day BID** | 4.2 ± 2.4 µg/ml<br>(8,4 ± 4,9 µM) | 3.7 ± 1.4 µg/g<br>(7,6 ± 2,9 µM) | 99 ± 27% |

\*: Trough concentrations
Data represent mean±SD (four animals per group)

## Discussion

Imatinib was previously identified and studied as an *in vitro* inhibitor of several coronaviruses in multiple cell-lines. It prevents the spike protein mediated fusion with the cell membrane by targeting the Abelson tyrosine-protein kinase 2 (Abl2) (Coleman et al., 2016; Dyall et al., 2014; Sisk et al., 2018). Our *in vitro* results confirm that this mode of action seems to be identical with the SARS-CoV-2. Indeed only Imatinib and Masitinib, that share the same targets (Abl 1, abl2 and c-KIT/CD117) (Dubreuil et al., 2009; Mahon et al., 2000), were able to inhibit the replication of SARS-CoV-2 *in vitro*. The third TKI tested, the Asciminib, which targets only Abl1 (Schoepfer et al., 2018), was found inactive. Altogether, these results suggest that SARS-CoV-2 also depends on Abl2 for its entry and replication in VeroE6 cells. Moreover a recent report also shown that Imatinib block SARS-CoV-2 entry using human pluripotent stem cell-derived lung and colonic organoids, with similar EC50, suggesting that this antiviral mechanism is conserved in multiple cell lines (Han et al., 2020).

However, in spite of its undeniable activity in several cell models, Imatinib was not active in bronchial human airway epithelia that had been reconstructed from human primary cells of bronchial biopsies, despite the expression of Abl2 in lung tissue (Uhlén et al., 2015). Imatinib also failed to reduce lung viral replication at the peak of replication. It is unlikely that an insufficient pulmonary diffusion of Imatinib may explain this result, since a good penetration of Imatinib in the pulmonary tract was observed with trough concentrations in lung tissue above the EC50 value with both dosing regimens tested. Another hypothesis may be the non-conservation of Abl2 kinase between hamsters and primates. However, it turns out that the gene coding for Abl2 is highly conserved among vertebrates (Colicelli, 2010). Another explanation may be a difference in term of viral spread/replication between a cell monolayer and more complex tissues such as lungs and HAE. Indeed, the proposed mode action of Imatinib is to interfere with the viral entry step, but this inhibition solely may be insufficient in a more complex three-dimensional tissue :once the virus has entered the cell, it may use another pathway to infect surrounding cells, like cell-to cell transfer as previously described for other enveloped human viruses (Merwaiss et al., 2019; Mothes et al., 2010; Sattentau, 2008).

Taken together, these results indicate that the antiviral activity of Imatinib is only present in cell cultures, and that its capacity to prevent viral replication is absent in a small animal model or in reconstructed bronchial human airway epithelia. These results highlight the need for combined approaches during the preclinical evaluation of antiviral compounds even in the case of developing a drug repurposing approach. Indeed, the use of relevant animal model and relevant tissue may give precious information to clinicians who would be able more to eliminate rapidly compounds without antiviral activity or that insufficiently diffused in targeted tissues. This have been already observed for the chloroquine where the molecule block efficiently the viral replication in veroE6 cells ((Liu et al., 2020; Wang et al., 2020) but displayed less potency in human lung cells (Hoffmann et al., 2020) and no activity in HAE and non-human primates (Maisonnasse et al., 2020). In conclusion, these results do not support the use of Imatinib and similar TKIs as antivirals for the treatment of Covid-19.

## Supporting information

Supplemental Figures revised

## Contributions

FT, AN, CS and XDL conceived the experiments. XDL, FXM and DM proposed the study. FT, JSD, MC, PRP, MG, KB, GM and AN performed the experiments. FT, JSD, MC, PRP, CS and AN analysed the results. FT, JSD, CS, AN and XDL wrote the paper. FT, JSD, MC, PRP, MG, KB, GM, FXM, DM, CS, XDL and AN reviewed and edited the paper.

## Acknowledgments

We would like to thank Camille Placidi-Italia for excellent technical support. We thank Alexandre Moussy from AB science for providing the Masitinib mesylate. We thank Marjorie Roudot from the hospital pharmacy of the University hospital of La Timone (Marseille) for providing 400mg tablets of Imatinib mesylate used for *in vivo* experiments. We thank FUJIFILM Toyama Chemical Co for kindly providing the Favipiravir. We thank Pr C Drosten for providing the SARS-CoV-2 strain through EVA GLOBAL. A part of the work was done on the Aix Marseille University antivirals drug design platform “AD2P”.

