## Supplemental Figures revised for "Preclinical evaluation of Imatinib does not support its use as an antiviral drug against SARS-CoV-2"

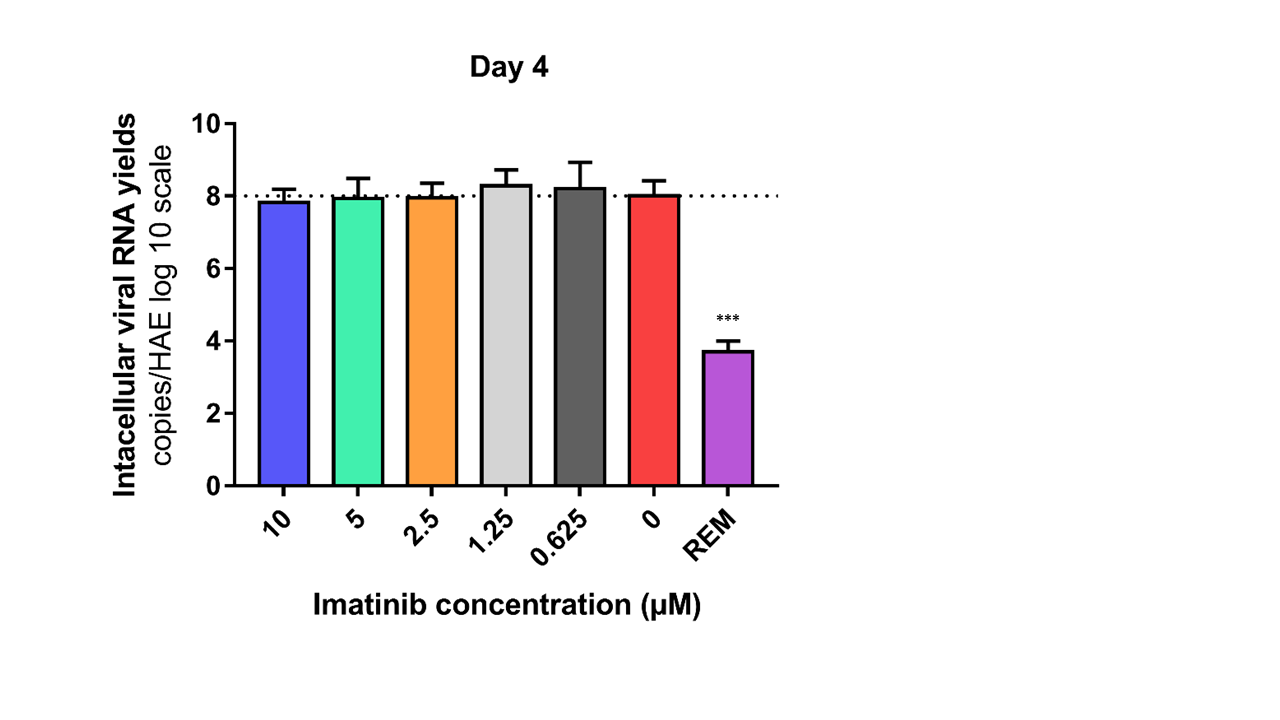

**Supplemental Figure 1: Intracellular SARS-CoV-2 viral RNA quantification at day 4 post infection**.

No statistical difference were observed concerning Imatinib. Statistical significance was calculated by 1- way ANOVA versus untreated group. REM: remdesivir was used as a positive drug control with a statistical difference compared to control group ***p value=0.0001

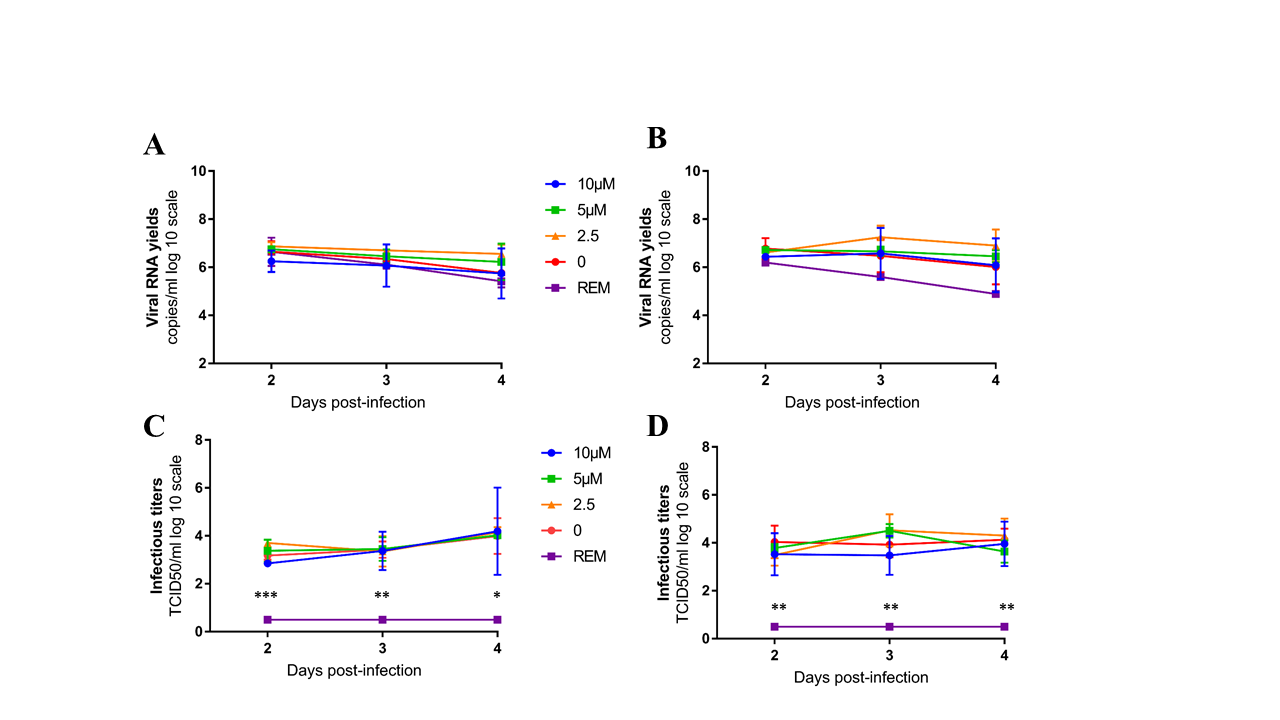

**Supplemental figure 2: Antiviral activity of Imatinib in a bronchial human airway epithelium**

Kinetics of virus excretion at the apical side of the epithelium measured using an RT-qPCR assay (A) and (B) and a TCID_50_ assay (C) AND (D) for confirmatory experiment 2 (A,C) and 3 (B,D) . Data represent mean±SD. Statistical significance was calculated by 1-way ANOVA versus untreated group. No statistical difference were observed with Imatinib regardless the drug conc. Rem (remdesivir at 10µM) was used as a positive drug control. *, ** and *** symbols indicate that the average value for the Rem group is significantly lower than that of the untreated group with a p-value ranging between 0.01-0.1, 0.001-0.01 and 0.0001-0.001 respectively.

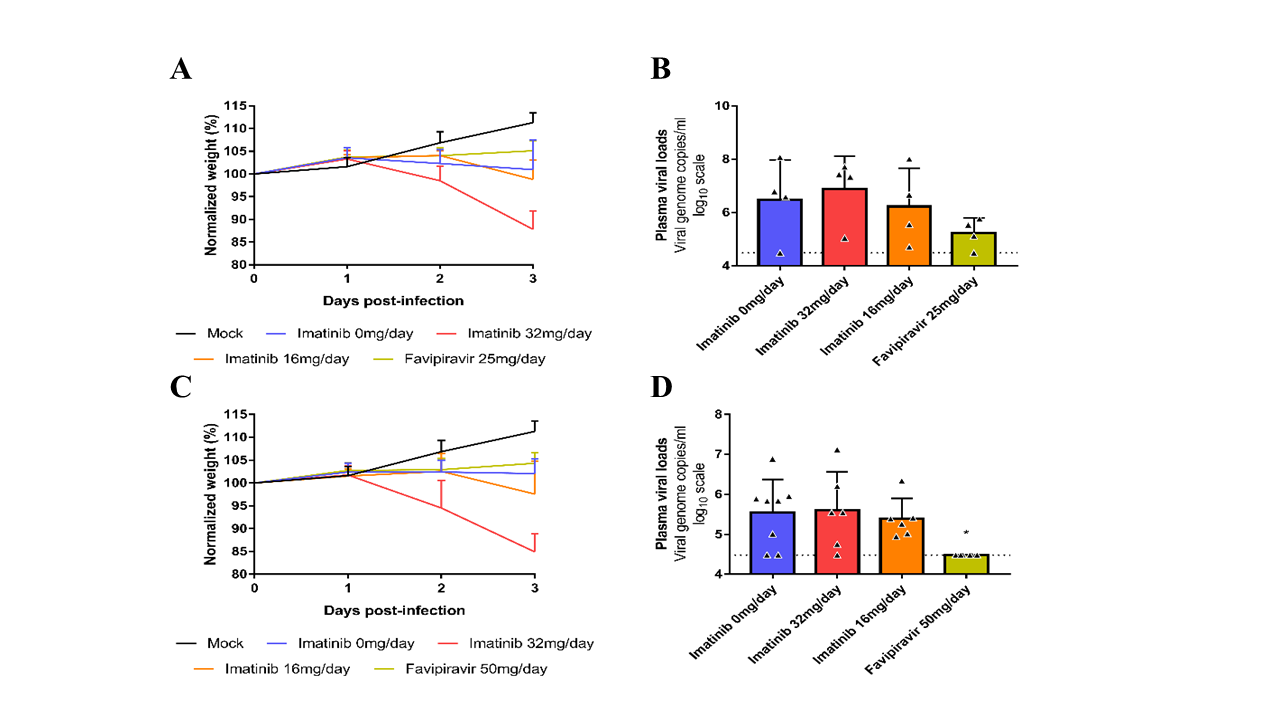

**Supplemental Figure 3**: **Clinica**l **follow-up of animals and plasma viral loads.**

A-B first experiment n=4, C-D second experiment n=6. A and C Treated and untreated animals normalized weight (% of initial weight of the animal at day n)/(mean % of initial weight for mock-infected animals at day n). B-D plasma viral load of animals (measured using RT-qPCR) expressed in viral genome copies/ml of plasma. Data represent mean ±SD.

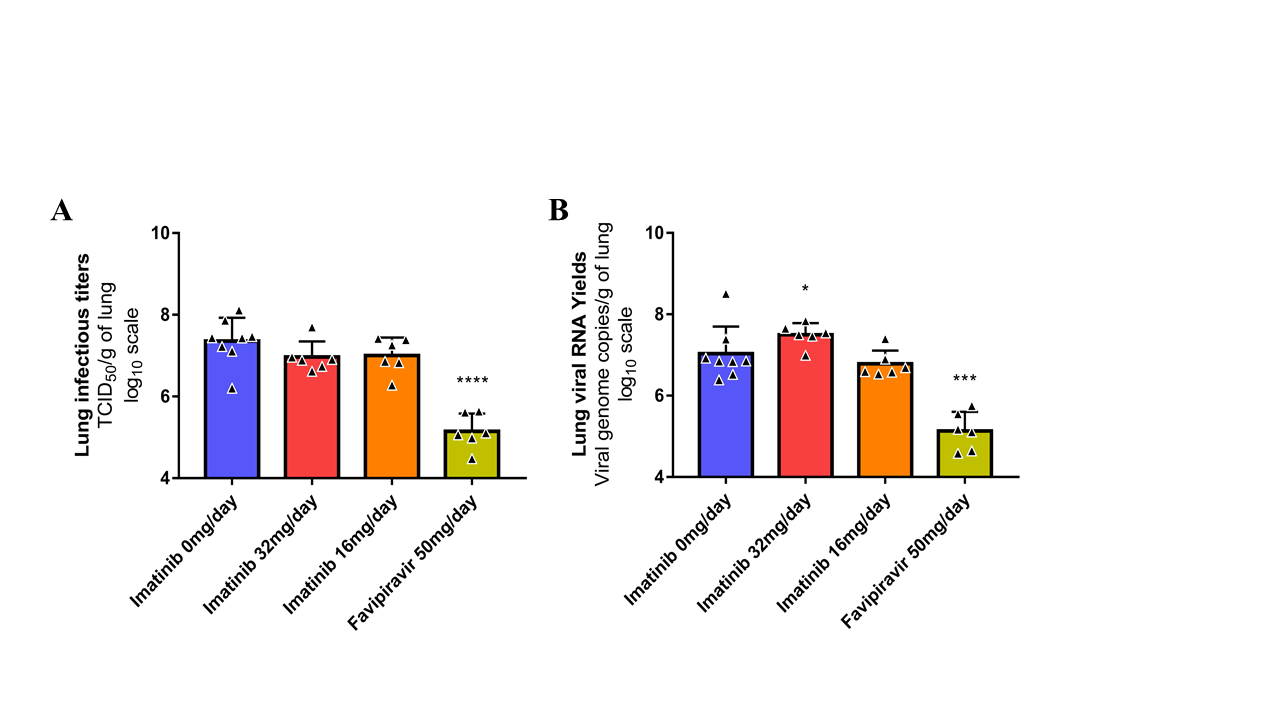

**Supplemental Figure 4:** **Antiviral activity of Imatinib in a Golden Syrian hamster model**:

Groups of 6 hamsters were intranasally infected with 10^4^ TCID_50_ of SARS-CoV-2 and received the Imatinib, orally twice a day. Favipiravir was used as a positive control. A: Lung infectious titers (measured using a TCID_50_ assay) expressed in TCID_50_/g of lung. B: Viral RNA yields (measured using an RT-qPCR assay) expressed in viral genome copies/g of lung. Data represent mean±SD. *** symbols indicate that the average value for the group is significantly lower than that of the untreated group with a p-value ranging between 0.0001-0.001.
